# Electrochemical Metabolic Profiling Reveals Mitochondrial Hyperactivation and Enhanced Neural Progenitor Proliferation in MLC1-Mutant Human Cortical Organoids

**DOI:** 10.1101/2025.04.16.647364

**Authors:** Kyeong-Mo Koo, Jeong-Sun Choi, Yun-Kyeong Nam, Hyung-Joo Kim, Chang-Dae Kim, Soo-Hwan Jeong, Hyun-Jong Jang, Kayeong Lim, Byung-Chan Lim, Tae-Hyung Kim, Kyung-Ok Cho

## Abstract

Mitochondrial function is critical for neural progenitor regulation, yet its dysregulation during early human brain development remains poorly defined. Megalencephalic leukoencephalopathy with subcortical cysts (MLC) is a neurodevelopmental disorder caused by MLC1 mutations, previously attributed to postnatal astrocyte dysfunction. Using patient-derived human cortical organoids, we show that MLC1 is expressed in early neuroepithelial cells. To assess mitochondrial state in live organoids, we developed the MAGO (Matrigel-coated gold nanostructure) platform for real-time, label-free detection of redox activity. MLC1 mutant organoids showed mitochondrial hyperactivation, increased ATP and ROS, reduced membrane potential, and altered fusion protein expression. These changes were accompanied by enhanced BrdU incorporation and expansion of PAX6⁺/SOX2⁺ progenitors. To assess the causal role of MLC1 mutation, we generated isogenic organoids using CRISPR prime editing, which recapitulated redox hyperactivation and increased proliferation. Our findings redefine MLC as a disorder of early mitochondrial and progenitor dysregulation and establish a tractable platform to study metabolic mechanisms in neurodevelopmental disease.

## INTRODUCTION

Mitochondrial metabolism plays a central role in regulating neural stem and progenitor cell behavior by coordinating the balance between proliferation and differentiation during early brain development^1–4^. These metabolic programs are closely coupled to developmental timing and lineage progression. Moreover, mitochondrial dysfunction has been implicated in several neurodevelopmental disorders (NDDs), including both primary mitochondrial diseases caused by mutations in mitochondrial genes and secondary dysfunctions arising from non-mitochondrial genetic etiologies^5–8^. However, despite accumulating evidence linking mitochondrial abnormalities to NDDs, current insights have mostly come from studies of mature neurons, and the specific contribution of mitochondrial dysregulation in human neural progenitor cells remains largely unexplored, partly due to limited access to early developmental stages *in vivo*.

Megalencephalic leukoencephalopathy with subcortical cysts (MLC) is a rare neurodevelopmental disorder primarily characterized by megalencephaly, white matter abnormalities, ataxia, seizures, and cognitive decline^9^. MLC is predominantly diagnosed within the first year of life, with symptoms progressively worsening over time The disorder is primarily caused by recessive mutations in *MLC1*, and less commonly by mutations in *GLIALCAM*, which can be inherited in either dominant or recessive forms^10,11^. The MLC1 gene, located on chromosome 22q13.33, consists of 12 exons and over 150 mutations have been identified across the MLC1 gene, which are predominantly associated with loss-of-function effects^12^. It is a devastating disease with no effective treatment currently available to cure or slow the disease progression. Therefore, it is essential to investigate the underlying molecular mechanisms to better understand disease pathogenesis and for enabling rational design of new diagnostic and therapeutic strategies.

MLC has traditionally been considered a disorder of astrocytic origin, based on the predominant expression of MLC1 in perivascular astrocytes^13,14^, where it is implicated in maintaining water and ion homeostasis^15^. Consequently, astrocyte dysfunction has been linked to hallmark features of MLC, including brain edema and white matter vacuolization. However, the diverse and progressive clinical manifestations of MLC, including cognitive impairment and motor dysfunction, suggest that astrocyte dysfunction alone may not fully explain the complex pathophysiology of the disease^9^. Recent technological advancements in human induced pluripotent stem cells (hiPSCs) reprogramming^16^ and CRISPR-Cas9 genome editing^17^ have enabled human patient-specific disease modeling and the introduction of precise MLC1 mutations in isogenic controls. Moreover, hiPSC-derived cortical organoid (hCO) models offer a powerful platform for investigating MLC pathologies by preserving the three-dimensional cellular architecture of the developing brain and providing an optimal model to study early neurodevelopmental alterations prior to the generation of astrocytes. Given that MLC1 is expressed in neuroepithelial cells in the early developing mouse brain^18^, establishing a robust cerebral organoid model for MLC will be crucial for elucidating its role in early cortical development and eventually developing novel therapeutic approaches for this devastating disorder.

In this study, we established a human cortical organoid (hCO) model of MLC using both patient-derived hiPSCs and prime-edited isogenic lines generated by introducing the same MLC1 mutation into wild-type hiPSCs. Both models exhibited heightened proliferative activity and significantly increased OXPHOS activity, as measured by our electrochemical (EC) platform, MAGO (MAtrigel-coated GOld nanostructured), which is a real-time, non-invasive EC system optimized for evaluating mitochondrial redox dynamics^19^. In patient-derived hCOs, this metabolic activation was accompanied by increased ATP and ROS levels, decreased mitochondrial membrane potential, and structural mitochondrial alterations, including elongation and selective changes in fusion regulators such as mitofusin 2 (MFN2) and short isoform of optic atrophy 1 (S-OPA1). Collectively, these findings uncover a previously unrecognized link between neural progenitor dysregulation and mitochondrial stress in MLC and establish a tractable model for probing early bioenergetic dysfunction in neurodevelopmental disorders.

## RESULTS

### MLC patient-derived hiPSCs maintain normal stem cell identity and multi-lineage potential

Peripheral blood mononuclear cells (PBMCs) from two MLC patients carrying a homozygous c.824C>A (p.Ala275Asp) mutation in MLC1^20^ were reprogrammed into human induced pluripotent stem cells (hiPSCs) using Sendai virus vectors encoding KLF4, OCT3/4, SOX2, and c-MYC. Sanger sequencing confirmed the presence of the same MLC1 mutation in hiPSCs as in the patients **(Figure 1A)**. Both control and mutant hiPSCs formed typical colonies and exhibited normal karyotypes **(Figures 1B and 1C)**. Quantitative PCR analysis showed significant upregulation of OCT4, SOX2, NANOG, and LIN28A mRNAs relative to PBMCs, with comparable expression between control and mutant lines **(Figure 1D)**. Immunostaining confirmed expression of OCT4, SOX2, NANOG, and SSEA-4 in both lines **(Figure 1E)**, confirming the maintenance of pluripotency in both control and MLC1 mutant hiPSCs. To evaluate differentiation potential, hiPSCs were subjected to trilineage induction. PAX6 (ectoderm), SM22A (mesoderm), and FOXA2 (endoderm) were robustly detected in both groups **(Figure 1F)**. These results confirm that MLC1 mutant hiPSCs retain pluripotency and trilineage competency.

**Figure 1.**
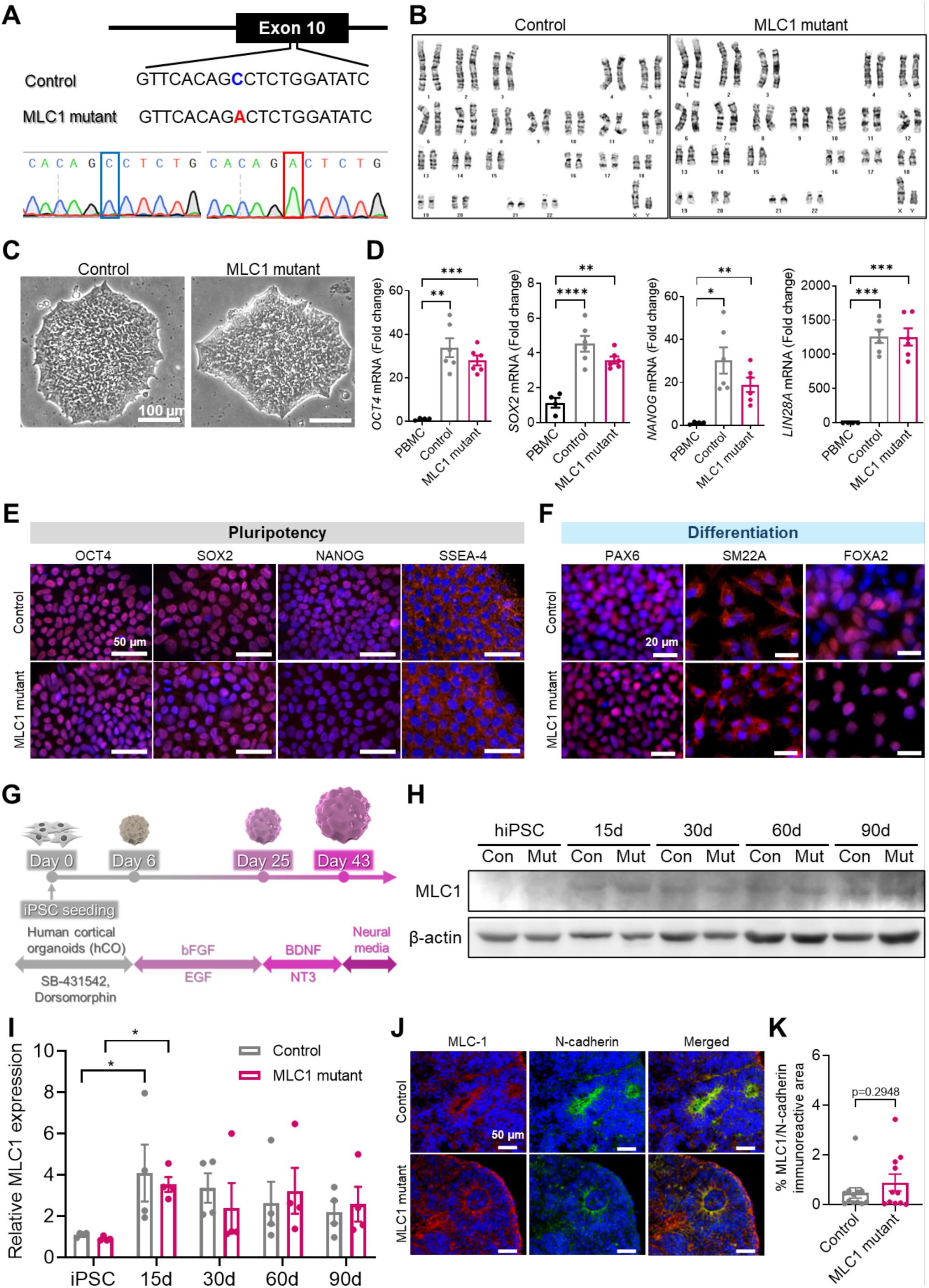
MLC1 patient-derived hiPSCs maintain pluripotency and hCOs express MLC1 during cortical development. (A) Sanger sequencing showing the homozygous c.824C>A (p.Ala275Asp) mutation in exon 10 of the MLC1 gene in patient-derived hiPSCs. (B) Karyotyping analysis confirming normal chromosomal integrity in both control and MLC1 mutant hiPSCs. (C) Representative bright-field images showing typical colony morphology of control and MLC1 mutant hiPSCs. (D) qPCR analysis showing comparable expression of pluripotency markers (OCT4, SOX2, NANOG, LIN28A) between control and MLC1 mutant lines (PBMC, n=4; Control, n=6; MLC1 mutant, n=6). (E) Immunofluorescence staining of OCT4, SOX2, NANOG, and SSEA-4 confirming pluripotency in both lines. Scale bars, 50 µm. (F) Immunostaining for PAX6 (ectoderm), SM22A (mesoderm), and FOXA2 (endoderm) in control and MLC1 mutant hiPSCs following directed differentiation. Scale bars, 20 µm. (G) Schematic showing human cortical organoid (hCO) generation from hiPSCs.(H) Western blot showing temporal expression of MLC1 protein in control and MLC1 mutant hCOs from day 15 to day 90. (I) Densitometric quantification of MLC1 protein levels normalized to β-actin, showing similar expression kinetics between groups (n=4). (J) Immunofluorescence of MLC1 and N-cadherin in day 30 hCOs. MLC1 is localized to neuroepithelial cells in both control and MLC1 mutant lines. Scale bar, 50 µm. (K) Quantification of MLC1⁺/N-cadherin⁺ immunoreactive area as a percentage of total hCO cross-sectional area, showing no significant difference between control (n=12) and MLC1 mutant hCOs (n=11). \**p* < 0.05, \*\**p* < 0.01, \*\*\**p* < 0.001, and \*\*\*\**p* < 0.0001. Data are presented as mean ± S.E.M.

### MLC1 is induced during early hCO development and remains unchanged by the patient mutation

hCOs were generated from control and MLC1 mutant hiPSCs using the Pasca protocol^21^ promoting dorsal forebrain specification **(Figure 1G)**. MLC1 protein was not detected in undifferentiated hiPSCs but peaked at day 15 in hCOs, followed by a gradual decline in both control and MLC1 mutant hCOs, as shown by Western blot **(Figures 1H and 1I)**. At day 30, MLC1 was localized to N-cadherin-expressing neuroepithelial cells in both control and mutant hCOs **(Figure 1J)**. Quantification of % MLC1⁺/N-cadherin⁺ immunoreactive area revealed no significant difference between groups **(Figure 1K)**. These results indicate that MLC1 is expressed during early cortical organoid development and that the patient-derived mutation does not alter its expression or localization in neuroepithelial cells.

### MAGO platform reveals persistent mitochondrial OXPHOS activation in MLC1 mutant hCOs

To enable quantitative monitoring of mitochondrial activity in hCOs, we developed an EC sensing platform—MAGO (MAtrigel-coated GOld nanostructured platform)—comprising nanostructured gold electrodes deposited on an ITO substrate **(Figure 2A)**. The morphological and topographical characteristics of MAGO were characterized using field-emission scanning electron microscopy (FE-SEM) and atomic force microscopy (AFM), confirming the formation of nanostructured surfaces with appropriate structural features **(Figures 2B and 2C)**. To minimize signal variation caused by surface heterogeneity, the gold nanostructured electrode was coated with Matrigel. AFM analysis revealed that Matrigel application significantly enhanced surface uniformity, reducing root-mean-square (RMS) roughness by 43.42% relative to the uncoated control. Importantly, Matrigel coating process did not compromise EC conductivity **(Figures 2D–F)**. This coating conferred two key advantages: improved adhesion of three-dimensional organoids to the electrode surface and reduced chip-to-chip variability, likely by minimizing nanoscale topographical inconsistencies. The stabilizing effect of Matrigel was further supported by the narrower distribution of EC signals in the coated group **(Figure 2G)**, emphasizing the importance of surface optimization for reliable organoid-based EC sensing.

**Figure 2.**
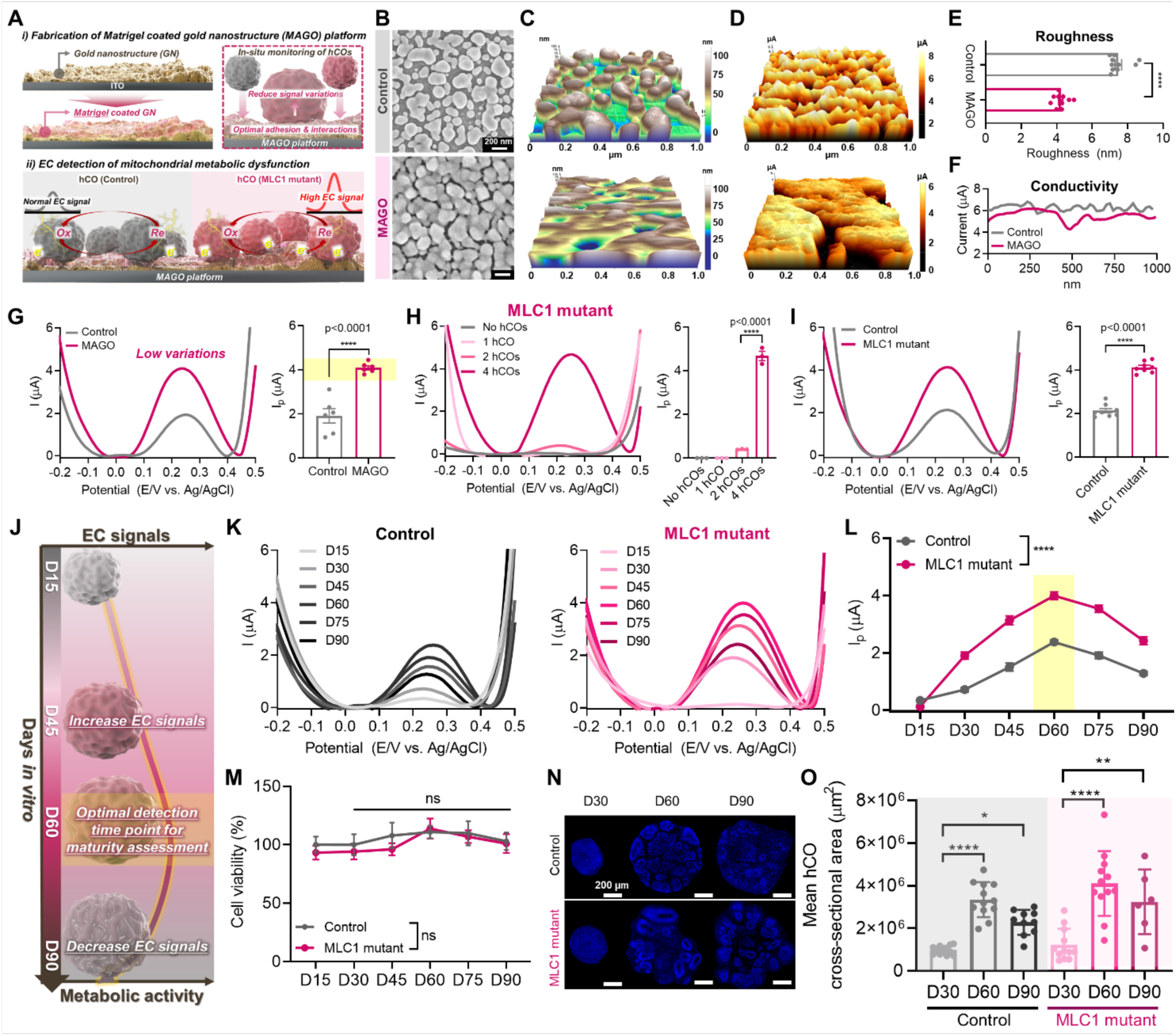
MAGO platform enables non-invasive temporal detection of mitochondrial OXPHOS activity, demonstrating persistent mitochondrial OXPHOS activation in MLC1 mutant hCOs. (A) Fabrication and EC detection of the MAGO (MAtrigel-coated GOld nanostructured) platform designed for *in situ* monitoring of mitochondrial activity in hCOs. (B-D) Characterization of the ITO/gold nanostructure (Control) and ITO/gold nanostructure/Matrigel (MAGO) platform using (B) FE-SEM images, (C) AFM analysis, and (D) conductive AFM (c-AFM) analysis. (E) Root mean square roughness (Rq) analysis for Control and MAGO platform groups (n=10). (F) Quantified results from the c-AFM images (D). (G) DPV graphs from MLC1 mutant organoids on Control and MAGO platform (left-panel) and the quantified DPV results presented as a bar graph (right-panel) (n=6). (H) DPV results with varying numbers of hCOs, ranging from 1 hCO to 4 hCOs (left-panel) and the quantified results presented as a bar graph (right-panel) (n=3). (I) DPV results from normal hCO (control) and MLC1 mutant hCO (left-panel) and the quantified results (n=7). (J) Scheme depicting the time-dependent monitoring of hCOs and MLC1 mutant models. (K) DPV graphs from Control (left-panel) and MLC1 mutant groups (right-panel) for 90 days. (L) Quantified DPV results presented as a line graph (n=5). (M) Cell viability tests after DPV detection presented as a line graph (n=5). (N) Representative DAPI images of control and MLC1 mutant hCOs at day 30, day 60, and day 90. Scale bar, 200 µm. (O) Mean hCO cross-sectional area of control and MLC1 mutant hCOs showing growth plateau beyond day 60, dissociating metabolic signals from physical expansion (Control: D30, n=12; D60, n=12, D90, n=10; MLC1 mutant: D30, n=12; D60, n=12, D90, n=6). \**p* < 0.05, \*\**p* < 0.01, \*\*\**p* < 0.001, and \*\*\*\**p* < 0.0001. Data are presented as mean ± S.E.M.

Next, we assessed the sensitivity of the MAGO platform by performing differential pulse voltammetry (DPV) with varying numbers of MLC1 mutant hCOs. As few as two organoids produced detectable signals, and increasing the number to four resulted in an 11.65-fold signal enhancement, demonstrating strong input-dependent amplification **(Figure 2H)**. This high sensitivity confirms that MAGO can detect mitochondrial metabolic activity with minimal sample input. We then compared metabolic activity between MLC1 mutant and control hCOs. MLC1 mutant hCOs exhibited a 1.94-fold increase in EC signal relative to controls **(Figure 2I)**. These results validate the utility of MAGO for detecting mutation-associated metabolic alterations in cortical organoid models.

To assess temporal changes in mitochondrial metabolic activity, we conducted serial measurements of hCOs using the MAGO platform from day 15 to day 90 at 15-day intervals **(Figures 2J and 2K)**. EC signals were minimal at day 15 but progressively increased in both control and MLC1 mutant hCOs, reaching a peak at day 60 **(Figure 2L)**. This signal elevation occurred in parallel with an increase in organoid cross-sectional area between day 30 and 60 **(Figures 2N and 2O)**, potentially related with developmental maturation and rising metabolic demand. From day 60 to 90, EC signals declined **(Figure 2L)** despite sustained size and viability **(Figures 2M–O)**, suggesting that the observed EC signals can reflect mitochondrial activity rather than total organoid mass or cell survival. Importantly, at all time points, MLC1 mutant hCOs consistently exhibited higher EC signals than controls **(Figure 2L)**. Together, these results validate the sensitivity of the MAGO platform for tracking temporal metabolic changes and reveal persistently elevated mitochondrial OXPHOS-associated signals in MLC1 mutant hCOs during early cortical development.

### MLC1 mutant hCOs exhibit mitochondrial hyperactivation, redox imbalance, and altered fusion dynamics

To elucidate the biochemical consequences of mitochondrial dysregulation observed in our MAGO platform, we conducted a series of functional and morphological assays at day 60. In galactose-based media to favor mitochondrial respiration, MLC1 mutant hCOs displayed significantly elevated basal ATP levels compared to controls, an effect abolished by oligomycin treatment, indicating enhanced OXPHOS-dependent ATP synthesis **(Figure 3A)**. Next, we examined mitochondrial membrane potential using FACS-based tetramethylrhodamine methyl ester (TMRM) fluorescence analysis. Under basal conditions, the mean TMRM fluorescence intensity was significantly reduced in MLC1 mutant hCO compared to controls, suggesting mitochondrial depolarization and potential mitochondrial dysfunction **(Figure 3B)**. This decrease in TMRM fluorescence was further confirmed upon treatment with the protonophore carbonyl cyanide m-chlorophenyl hydrazone (CCCP), which completely dissipated the mitochondrial membrane potential in both control and MLC1 mutant hCO **(Figure 3B)**.

**Figure 3.**
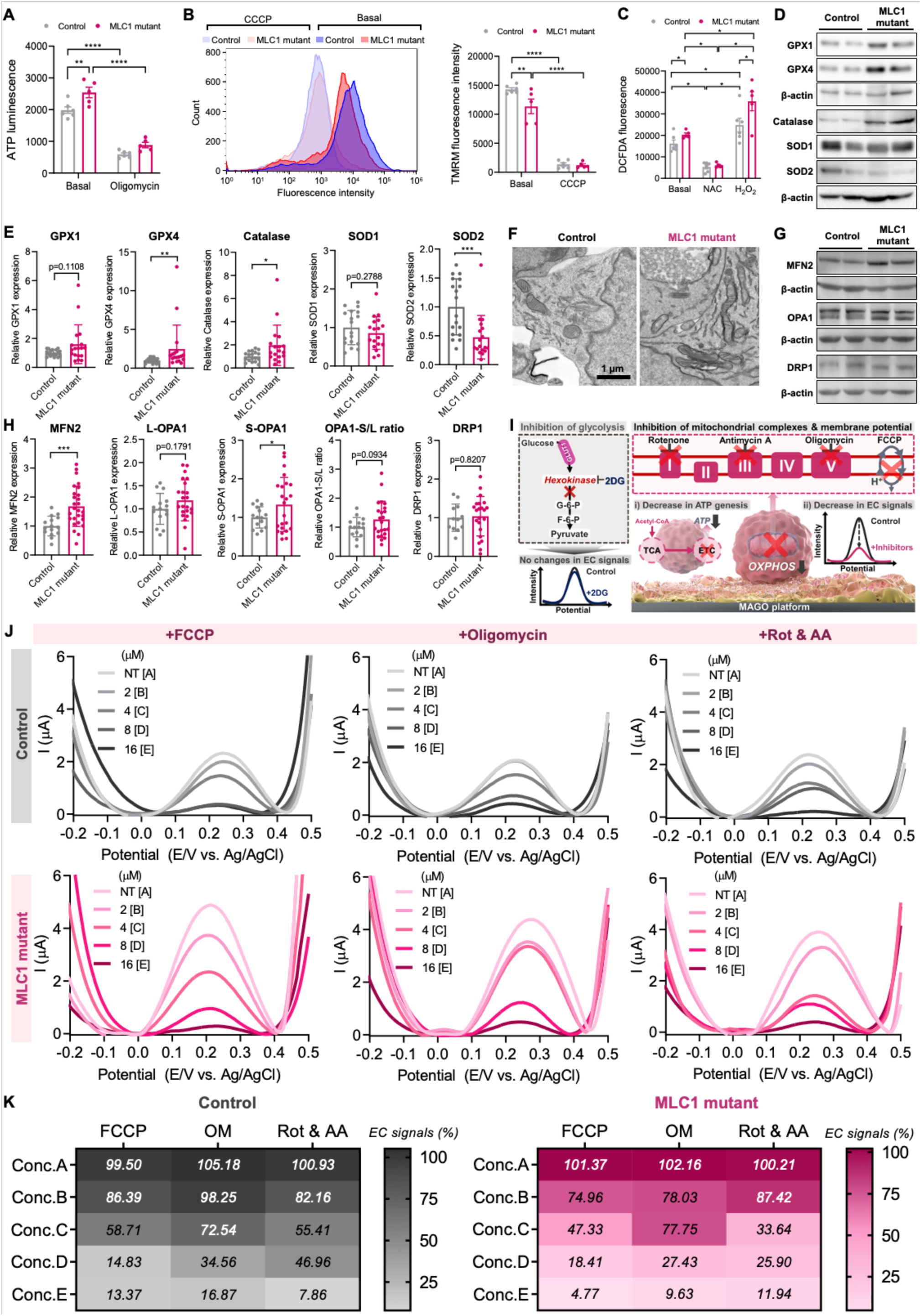
MLC1 mutant hCOs exhibit enhanced ATP levels, oxidative stress, altered fusion dynamics, and enhanced vulnerability to mitochondrial perturbation. (A) Intracellular ATP levels measured under galactose-based conditions at day 60. MLC1 mutant hCOs show elevated ATP levels, which are suppressed by oligomycin, indicating enhanced OXPHOS-dependent ATP production (Control, n=6; MLC1 mutant, n=5). (B) Mitochondrial membrane potential assessed via TMRM fluorescence by flow cytometry. Basal TMRM signal is reduced in MLC1 mutant hCOs and completely abolished by CCCP (Control, n=6; MLC1 mutant, n=5). (C) Intracellular ROS levels measured using H₂DCFDA fluorescence. MLC1 mutant hCOs display elevated basal ROS levels, which are attenuated by NAC and enhanced by H₂O₂ (Control, n=6; MLC1 mutant, n=5). (D) Western blot analysis of antioxidant enzymes. GPX4 and catalase are upregulated, while SOD2 is downregulated in MLC1 mutant hCOs relative to controls. (E) Densitometric quantification of protein expression from (D), normalized to β-actin (n=18). (F) Transmission electron microscopy showing elongated mitochondrial morphology in MLC1 mutant hCOs. Scale bar, 1 µm. (G) Western blot of mitochondrial fusion/fission regulators. MFN2 and S-OPA1 are increased in MLC1 mutant hCOs, while L-OPA1 and DRP1 remain unchanged. (H) Quantification of mitochondrial dynamics markers and OPA1 S/L ratio from (G) (MFN2: Control, n=15; MLC1 mutant, n=27, OPA-1: Control, n=15; MLC1 mutant, n=23, DRP1: Control, n=12; MLC1 mutant, n=22). (I) Schematic of EC detection in control and MLC1 mutant hCOs treated with glycolytic and mitochondrial inhibitors. (J) DPV graphs of hCOs and MLC1 mutant hCOs after 24-hour exposure to various concentrations of FCCP, oligomycin, and rotenone & antimycin A. (K) Heatmap depicting electrical signals in hCOs and MLC1 mutant hCOs (n=3) following the administration of mitochondrial inhibitors, derived from DPV graph (J). \**p* < 0.05, \*\**p* < 0.01, \*\*\**p* < 0.001, and \*\*\*\**p* < 0.0001. Data are presented as mean ± S.E.M.

To assess whether this bioenergetic imbalance was associated with oxidative stress, we measured intracellular ROS levels using H₂DCFDA fluorescence analysis. ELISA results revealed a significant increase in basal H_2_DCFDA fluorescence intensity in MLC1 mutant hCO compared to controls, indicating elevated ROS levels **(Figure 3C)**. Moreover, treatment with 1 mM N-acetyl-L-cysteine (NAC) significantly reduced H_2_DCFDA fluorescence intensity in both control and MLC1 mutant hCO. In contrast, administration of 2 mM hydrogen peroxide (H_2_O_2_) increased H_2_DCFDA fluorescence intensity in both groups compared to basal levels, with a significantly greater increase observed in MLC1 mutant hCO than controls. These findings suggest that despite enhanced OXPHOS activity, MLC1 mutant hCO are subjected to increased oxidative stress, which may contribute to mitochondrial dysfunction. To further investigate oxidative stress responses, we assessed the expression levels of key antioxidant enzymes such as GPX1, GPX4, catalase, SOD1, and SOD2 via Western blot analysis. Compared to control hCO, GPX4 and catalase were significantly upregulated in MLC1 mutant hCO, suggesting a compensatory response to elevated oxidative stress **(Figures 3D and 3E)**. Intriguingly, SOD2 was significantly downregulated in MLC1 mutant hCO compared to controls **(Figures 3D and 3E)**, implying that increased oxidative stress in MLC1 mutant hCO may be attributed from a reduced capability to convert mitochondrial superoxide anions into H_2_O_2_.

To determine whether these metabolic abnormalities were associated with mitochondrial structural remodeling, we analyzed mitochondrial morphology and dynamics regulators. Electron microscopy revealed elongated mitochondrial morphology in MLC1 mutant hCOs relative to controls **(Figure 3F)**. In support of our electron microscopic findings, Western blot analysis demonstrated that MFN2 expression, which mediates outer membrane fusion, was significantly increased, while levels of DRP1 remained unchanged **(Figures 3G and 3H)**. The short isoform of OPA1 (S-OPA1) was significantly upregulated in the absence of changes in the long isoform (L-OPA1), suggesting proteolytic processing in response to mitochondrial stress **(Figures 3G and 3H)**. Together, these findings indicate that MLC1 mutation leads to sustained OXPHOS activity, redox imbalance, and abnormal mitochondrial fusion, contributing to a metabolically overactive yet structurally imbalanced mitochondrial phenotype in early cortical development.

### Pharmacological inhibition of mitochondrial complexes and membrane potential validates OXPHOS specificity of MAGO platform-based EC signals in MLC1 mutant hCO

To further validate that EC signals predominantly originate from mitochondrial redox activity, we performed pharmacological metabolic perturbation experiments using our previous validated EC platform **(Figure 3I)**, which links cellular metabolic state and redox dynamics^19^. Following treatment with 2-deoxy-D-glucose (2DG), a glycolytic inhibitor targeting hexokinase, no significant changes in EC signal were observed in either control or MLC1 mutant hCOs **(Figure S2)**, indicating that glycolysis contributes minimally to the EC signals under these conditions. In contrast, treatment with mitochondrial inhibitors—including rotenone (complex I), antimycin A (complex III), oligomycin (complex V), and the uncoupler carbonyl cyanide-p-trifluoromethoxyphenylhydrazone (FCCP)—induced a robust reduction in EC signals in both groups **(Figure 3J)**, substantiating that the measured signals predominantly originate from mitochondrial activity. Notably, MLC1 mutant hCOs showed a markedly enhanced sensitivity of EC signals to mitochondrial perturbation **(Figure 3K)**. Specifically, at low concentration of FCCP (2 µM; Conc.B), EC signal reduction was 11.43% greater in mutants than in controls. Oligomycin (2 µM) produced an even larger differential, with a 20.22% stronger decrease in the mutant group. Furthermore, combined inhibition of complex I and III with rotenone and antimycin A (4 µM) induced a dramatic 66.36% signal suppression in MLC1 mutant hCOs, highlighting a heightened vulnerability in mitochondrial electron transport activity and bioenergetic resilience.

### Increased proliferative activity in both patient-derived MLC1 mutant hCO and prime-edited MLC1 mutant hCO

To explore the cellular basis of enhanced mitochondrial OXPHOS in MLC1 mutant hCOs, we performed immunostaining for PAX6 and SOX2 to label radial glia and neural progenitors. In both control and MLC1 mutant hCOs at day 30 and day 60, PAX6⁺ and SOX2⁺ cells were localized predominantly in rosette-like radial structures resembling the ventricular zone (VZ) in developing cortex **(Figures 4A and 4B)**. Quantitative analysis showed a significant increase of % PAX6- and SOX2-immunoreactive areas in MLC1 mutant hCOs compared to controls at days 30 and 60 **(Figures 4C and 4D)**. To directly assess whether this expansion reflected increased proliferative activity, we performed a BrdU pulse-labeling experiment. hCOs were treated with 10 µM BrdU for 24 h at day 30 and analyzed 7 days later. BrdU+ cells were localized predominantly around the rosette structures, and their area was significantly greater in MLC1 mutant hCOs than in controls **(Figure 4E)**. These findings indicate that enhanced stem/progenitor expansion and proliferation underlie the elevated metabolic demand observed in MLC1 mutant hCOs.

**Figure 4.**
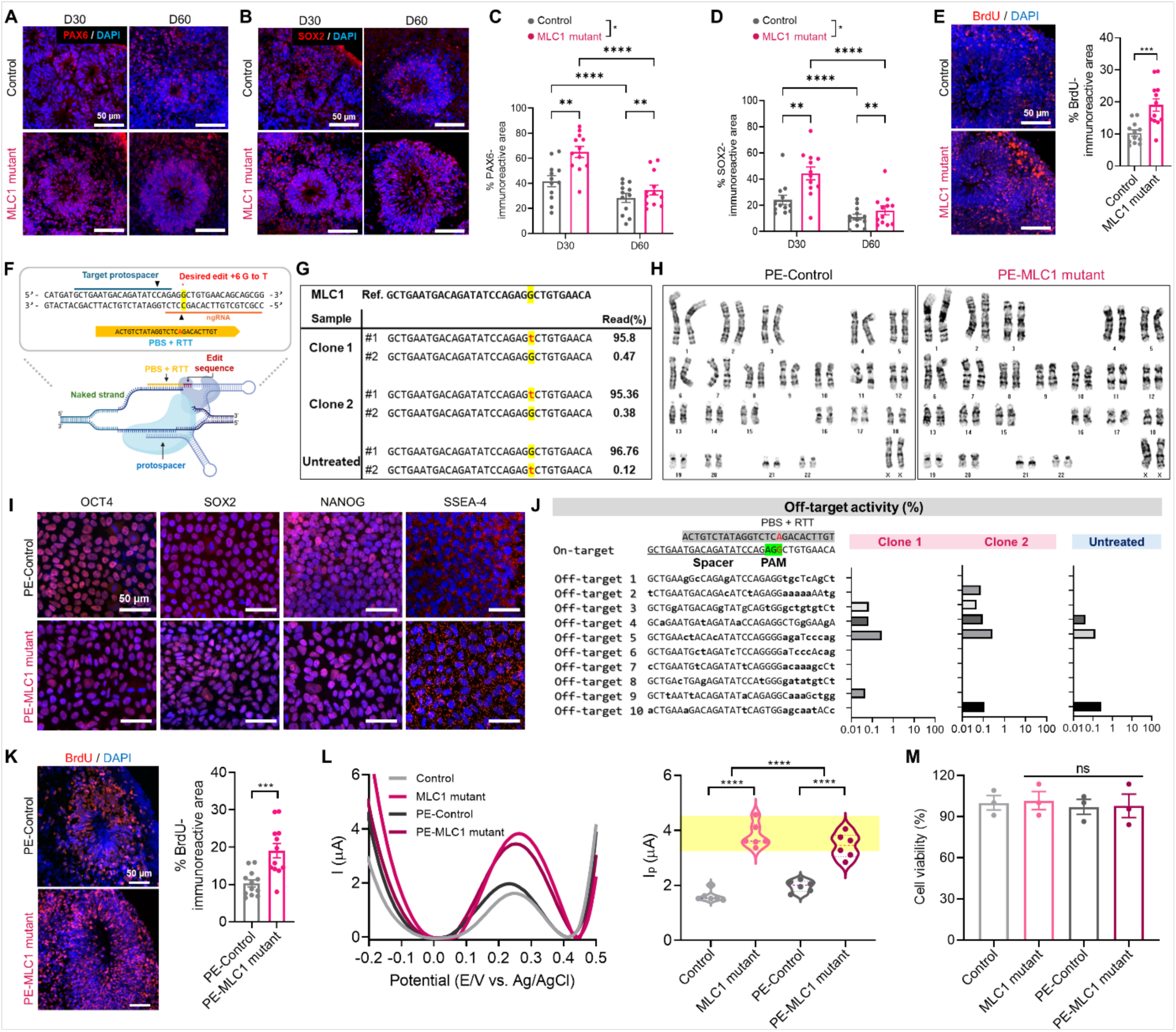
Increased proliferative activity and metabolic hyperactivation in both patient-derived and prime-edited MLC1 mutant hCOs. (A, B) Immunofluorescence staining of PAX6 (A) and SOX2 (B) in control and MLC1 mutant hCOs at days 30 and 60, showing enriched expression in rosette structures. Scale bars, 50 µm. (C, D) Quantification of PAX6⁺ and SOX2⁺ immunoreactive areas at days 30 and 60. MLC1 mutant hCOs show significantly expanded progenitor populations (n=12). (E) BrdU pulse-chase assay (24 h pulse on day 30, analysis on day 37) showing increased BrdU⁺ area in MLC1 mutant hCOs (n=12). Scale bar, 50 µm. (F) Schematic of CRISPR-mediated prime editing to introduce the c.824C>A (p.Ala275Asp) mutation into control hiPSCs. (G) Next-generation sequencing confirms >95% biallelic editing in two selected prime editing-derived clones. (H) Representative karyotyping result showing normal chromosomal integrity in prime-edited MLC1 mutant hiPSCs. (I) Immunostaining of OCT4, SOX2, NANOG, and SSEA-4 confirming maintenance of pluripotency in prime-edited lines. Scale bar, 50 µm. (J) Targeted deep sequencing of top 10 predicted off-target sites showing no detectable off-target editing in prime-edited clones compared to untreated controls. (K) BrdU pulse-chase assay (24 h pulse on day 30, analysis on day 37) showing increased BrdU⁺ area in prime-edited MLC1 mutant hCOs (PE-Control, n=5; PE-MLC1 mutant, n=4). Scale bar, 50 µm. (L) DPV graphs of hCOs, MLC1 mutant hCOs, PE-Control hCOs, and PE-MLC1 mutant hCOs (left-panel) and the quantified DPV results (right-panel) (n=6). (M) Cell viability tests after DPV detection are presented as a bar graph (n=3). \**p* < 0.05, \*\**p* < 0.01, \*\*\**p* < 0.001, and \*\*\*\**p* < 0.0001. Data are presented as mean ± S.E.M.

To determine whether the MLC1 mutation alone is sufficient to elicit the proliferative and metabolic phenotypes observed in patient-derived hCOs, we introduced the c.824C>A (p.Ala275Asp) point mutation into control hiPSCs using prime editing (PE) **(Figure 4F)**. Next-generation sequencing of clonal lines confirmed >95% editing efficiency for both selected clones **(Figure 4G)**, indicating successful editing in both alleles. Karyotype analysis showed normal chromosomal integrity in the prime-edited MLC1 mutant hiPSCs **(Figure 4H)**, and immunostaining verified that prime-edited MLC1 mutant hiPSCs maintained expression of pluripotency markers such as OCT4, SOX2, NANOG, and SSEA-4 **(Figure 4I)**. To assess potential off-target effects, we performed targeted deep sequencing of the top ten predicted sites. No off-target editing above background levels was detected in two clones compared to untreated controls **(Figure 4J)**. These results confirm the generation of isogenic MLC1 mutant hiPSC lines with minimal off-target risk and preserved pluripotency.

Next, we generated hCOs using prime-edited MLC1 mutant hiPSCs and evaluated their proliferative and metabolic phenotypes. Consistent with the patient-derived hCOs, prime-edited MLC1 mutant hCOs exhibited significantly increased BrdU incorporation at day 37 compared to isogenic controls **(Figure 4K)**. Electrochemical analysis using the MAGO platform also revealed elevated OXPHOS-associated EC signals in prime-edited mutant hCOs, recapitulating the metabolic hyperactivity observed in patient-derived models **(Figure 4L)**. Importantly, these changes occurred in the absence of differences in cell viability **(Figure 4M)**, supporting the conclusion that MLC1 mutation is sufficient to induce both excessive progenitor proliferation and mitochondrial metabolic activation, underscoring a causal link between MLC1 mutation and early neurodevelopmental dysregulation.

## DISCUSSION

MLC1 has primarily been studied in the context of postnatal astrocyte function^13,14,22^, and its role during earlier stages of brain development remains poorly understood. Investigating this question has been complicated by the cellular complexity of *in vivo* systems, where neural stem cells differentiate in parallel with the emergence of endothelial cells, microglia, and oligodendrocytes. Human cortical organoids (hCOs), which recapitulate early corticogenesis through the stepwise generation of neural stem, progenitor, and astrocyte lineages while lacking immune, vascular, and oligodendrocyte populations^21,23,24^, provide a controlled model to examine lineage-intrinsic roles of MLC1 with temporal precision. Using both patient-derived and prime-edited MLC1 mutant hCOs, we observed that MLC1 mutation disrupts neural stem/progenitor proliferation and mitochondrial metabolic homeostasis as early as day 30 of development. At this stage, MLC1 protein was localized in N-cadherin+ neuroepithelial cells within rosette structures. In addition, MLC1 protein expression was absent in hiPSCs, became strongly induced at day 15 of hCO development, and gradually declined thereafter in both control and MLC1 mutant hCO. The presence of MLC1 in these early neural compartments likely underlies the defects observed prior to astrocyte emergence. These findings suggest that MLC1 contributes to early cortical development independently of astrocytic context. Supporting this, *in situ* hybridization in embryonic mouse brains revealed Mlc1 mRNA expression in the ventricular zone at E18, which declined postnatally^25^. An Mlc1-T2A-eGFP reporter mouse also showed promoter activity in neuroepithelial cells at E14, prior to the onset of astrocyte differentiation^18^. Together, these results define a previously unrecognized role for MLC1 during early corticogenesis and underscore the utility of hCOs as a platform to dissect stage-specific functions of genes implicated in neurodevelopmental disease.

Our electrochemical analysis with MAGO platform during early cortical development revealed a dynamic trajectory of mitochondrial activity in hCOs. In control hCOs, OXPHOS activity showed minimal levels at day 15, peaked at day 60, and declined by day 90. In previous studies, hiPSC-derived neural systems have been shown to exhibit a glycolytic metabolic bias during early stages, with a gradual transition to oxidative phosphorylation as differentiation progresses^26,27^, suggesting that the initial increase in OXPHOS activity observed in our control hCOs reflects maturation-associated metabolic reprogramming. Notably, OXPHOS signals remained significantly elevated in MLC1 mutant hCOs at all time points, indicating a sustained enhancement of mitochondrial respiration that is independent of glycolysis, as evidenced by its insensitivity to 2-deoxyglucose and suppression by mitochondrial complex inhibitors. At day 60, despite a reduction in mitochondrial membrane potential and increased ROS levels, mutant hCOs produced more ATP under galactose medium. This paradoxical increase in ATP production, along with preserved ATP synthase coupling as shown by oligomycin sensitivity and responses to FCCP/CCCP, suggests a high-flux, low-potential mode of OXPHOS that may arise from enhanced electron transport chain activity under altered membrane conditions. At the structural level, our electron microscopic analysis demonstrated elongated mitochondrial morphology in MLC1 mutant hCOs, compared to more rounded morphology in controls. Western blot analysis showed a significant increase in MFN2 and the short isoform of OPA1 (S-OPA1) without corresponding changes in the long isoform (L-OPA1) or DRP1. Given that S-OPA1 is typically produced by proteolytic cleavage of L-OPA1 under stress conditions, this selective increase aligns with our observations of decreased mitochondrial membrane potential, elevated ROS, and reduced SOD2 expression. Although these findings indicate an enhancement of outer membrane fusion, inner membrane fusion or cristae remodeling may not be proportionally activated. Such uncoupled dynamics could contribute to the bioenergetic inefficiency and oxidative stress observed in MLC1 mutant hCOs, as shown in previous studies^28–31^. Together, these findings suggest that MLC1 mutation induces a metabolically overactive yet structurally imbalanced mitochondrial phenotype, linking altered fusion dynamics and redox dysregulation to early-stage developmental misregulation.

We next examined whether increased energy demand was associated with altered cellular proliferation and found significantly increased BrdU incorporation in MLC1 mutant hCOs as early as day 30, accompanied by an expansion of PAX6+ and SOX2+ progenitor populations. This elevated energy requirement from actively proliferating progenitors may contribute to mitochondrial hyperactivity, leading to bioenergetic imbalance and oxidative stress, ultimately resulting in mitochondrial dysfunction. As loss of mitochondrial fusion proteins such as MFN2 or OPA1 has been shown to reduce neural progenitor proliferation and self-renewal^2^, the concurrent increase in MFN2 expression and progenitor proliferation in our MLC1 mutant hCOs is consistent with this relationship. While mitochondrial dysfunction has been implicated in various neurodevelopmental disorders, including autism spectrum disorder and intellectual disability^5,6,30^, these are typically characterized by reduced OXPHOS activity and diminished ATP production. In contrast, MLC1 mutant hCOs showed a distinct hypermetabolic profile that may reflect a maladaptive cellular state arising from sustained proliferative pressure. These findings reveal a novel pathophysiological mechanism in MLC, diverging from previously reported mitochondrial impairments, and highlighting the complex interplay between proliferation and metabolism during early cortical development.

To establish a direct causal link between MLC1 mutation and the observed phenotypes, we introduced a patient-specific MLC1 mutation into control hiPSCs using prime editing. After confirming the absence of off-target effects, we successfully generated isogenic hiPSC lines harboring biallelic MLC1 mutations. These prime-edited mutant hCOs exhibited significantly increased BrdU incorporation at 37 days of development and elevated EC signals, recapitulating the enhanced proliferation and mitochondrial activation observed in patient-derived hCOs. These findings demonstrate that MLC1 mutation alone is sufficient to drive the neurodevelopmental phenotypes identified in our patient-derived hCO model and confirm its pathogenic role independent of other genetic backgrounds.

Although MLC is clinically characterized by early-onset macrocephaly and developmental delay, the dysregulation of progenitor proliferation and metabolic stress observed in our study may subtly disrupt cortical development and contribute to long-term functional outcomes such as cognitive impairment. Importantly, these early cellular changes occur at a stage that is not captured by standard clinical assessments such as MRI. While genetic testing can identify MLC1 mutations, it provides no information about the onset or trajectory of cellular dysfunction. This highlights a critical gap in current diagnostic approaches, emphasizing the need for tools that can capture early-stage pathophysiological changes. Our MAGO platform offers a powerful approach to address this cap by enabling sensitive, *in situ* assessment of mitochondrial OXPHOS activity across developmental time. By integrating nanoscale surface engineering with Matrigel-coated interfacing, this system allows for sensitive detection of mitochondrial activity, discrimination of disease-associated metabolic phenotypes, and temporal tracking of bioenergetic dynamics during early cortical development. Furthermore, these findings reinforce that EC signals in 3D brain organoid systems predominantly reflect mitochondrial function and, more importantly, uncover a previously unrecognized metabolic vulnerability in MLC1 mutant hCOs. The MAGO platform thus provides both mechanistic insight into early-stage metabolic dysregulation and a non-invasive framework for evaluating mitochondrial health in human neural models. These results support the application of EC-based sensing as a complementary tool to genetic testing, with potential utility for early detection of bioenergetic dysfunction and for stratifying patients for future therapeutic interventions in neurodevelopmental disorders such as MLC.

In summary, our findings demonstrate that MLC1 mutation induces a coordinated shift in neural progenitor behavior and mitochondrial metabolism during early corticogenesis. Both patient-derived and prime-edited MLC1 mutant hCOs exhibited enhanced cellular proliferation accompanied by increased oxidative phosphorylation activity. In patient-derived hCOs, this metabolic shift was further associated with elevated ATP production, increased reactive oxygen species likely resulting from reduced SOD2 expression, and impaired mitochondrial membrane potential. Mitochondrial elongation and selective MFN2 upregulation further support altered fusion dynamics that may contribute to metabolic activation. Taken all together, our results demonstrate a dual phenotype of hyperproliferation and mitochondrial stress in MLC1 mutant hCOs, providing insight into how metabolic dysregulation may impair early cortical development in MLC. Moreover, our MAGO platform successfully captured these early bioenergetic abnormalities in both patient-derived and prime-edited MLC1 mutant hCOs, supporting its potential as a diagnostic platform for identifying disease-relevant metabolic alterations during early cortical development.

## RESOURCE AVAILABILITY

### Lead contact

Kyung-Ok Cho is the lead contact for this manuscript and should be contacted for further information and requests for resources and reagents.

### Materials availability

This study did not generate new unique reagents. And Source data can be found in Data S1.

### Data and code availability

An Excel file containing the values to create graphs in the paper. This paper did not report the original code. Any additional information required to reanalyze the data reported in this paper is available from the lead contact upon request.

## ACKNOWLEDGEMENTS

This research was supported by the Nano & Material Technology Development Program through the National Research Foundation of Korea (NRF) grant funded by the Korea government, the Ministry of Science and ICT (MSIT) (Grant No. RS-2024-00410437 and RS-2024-00342716), and a grant of the Korea Health Technology R&D Project through the Korea Health Industry Development Institute (KHIDI), funded by the Ministry of Health & Welfare, Republic of Korea (RS-2023-00265923) This research was also supported by Basic Medical Science Facilitation Program through the Catholic Medical Center of the Catholic University of Korea funded by the Catholic Education Foundation, and by the Korea Institute of Science and Technology (KIST) Institutional Program (2V10573 and 2E33791). We thank Ms. Hyojung Lee and Ms. Yun-Mi Kim for their technical support. We also thank Dr. Jiyoon Kim (The Catholic University of Korea) for providing his electroporator and Dr. Eun-Kyung Lee (The Catholic University of Korea) for her valuable comments on mitochondrial dynamics.

## AUTHOR CONTRIBUTIONS

T.-H.K. and K.-O.C. supervised the project and conceived the idea. K.-M.K. and J.-S.C. contributed equally to this project. K.-M.K. and J.-S.C. sourced materials and conducted experiments. K.-M.K. H.-J.K., and C.-D.K. developed the MAGO platform for monitoring hCOs using the EC method. K.-O.C. generated hiPSCs and hCOs. K.-M.K. carried out most of the EC detection of hCOs under T.-H.K.’s supervision. J.-S.C. performed all the histology and Y.-K.N. carried out western blot analysis. H.-J.J. developed an automated imaging processing algorithm. K.L. and S.-H.J. designed plasmid constructs for prime editing the MLC1 locus and validated the prime-edited MLC1 mutant hiPSCs. B.-C.L. acquired patient samples and performed validation experiments. K.-M.K., J.-S.C., B.-C.L., T.-H.K., and K.-O.C. analyzed and interpreted the data.

## STAR★METHODS

### KEY RESOURCES TABLE

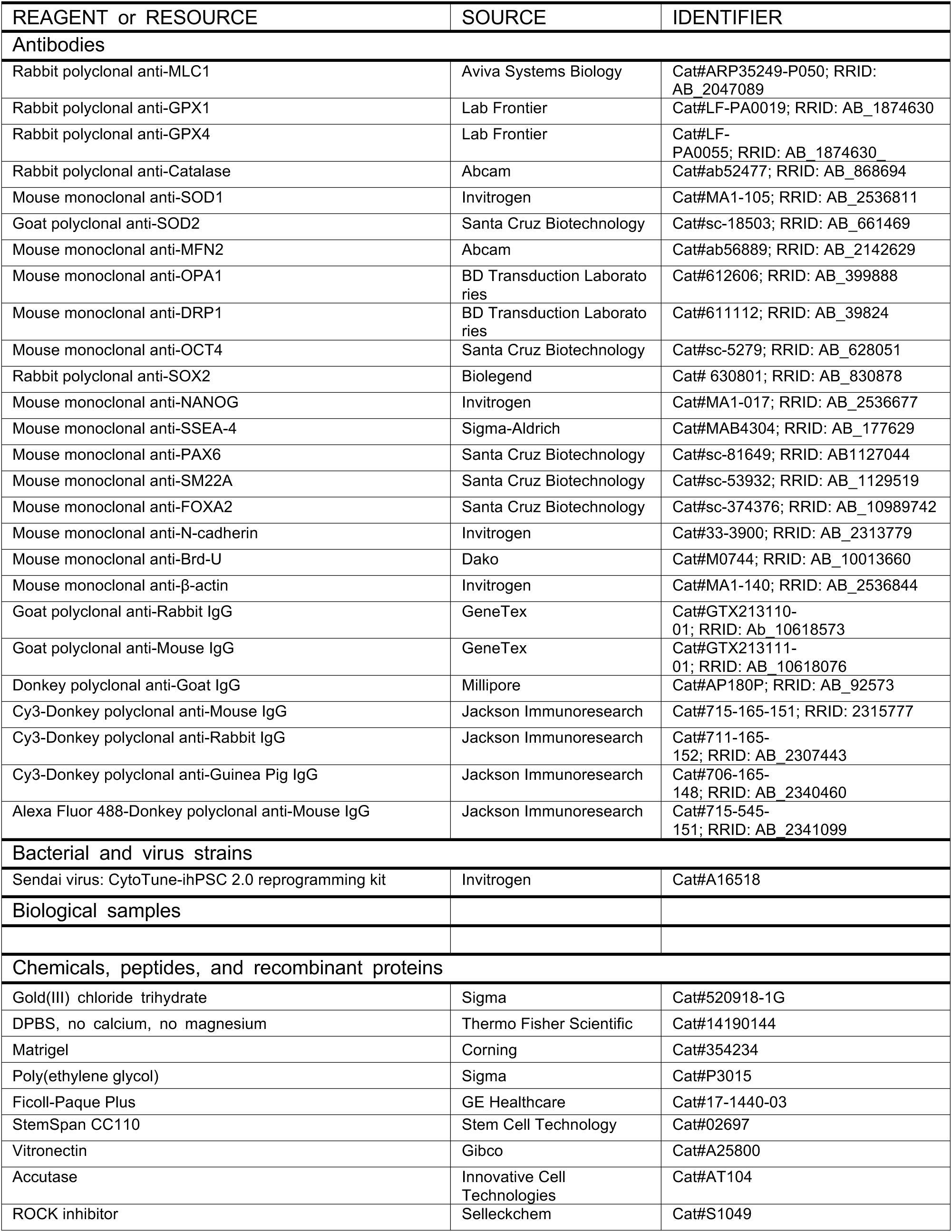
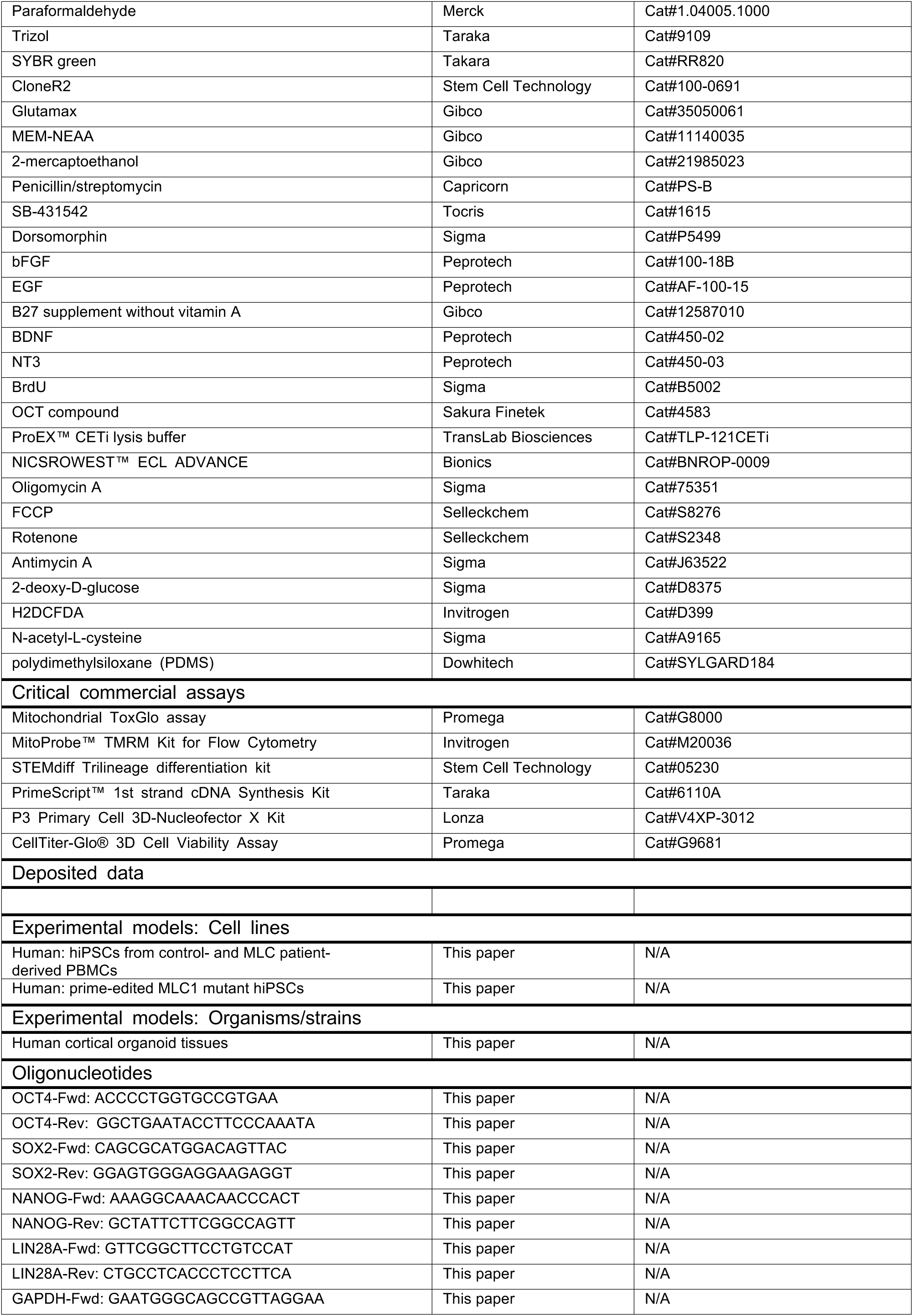
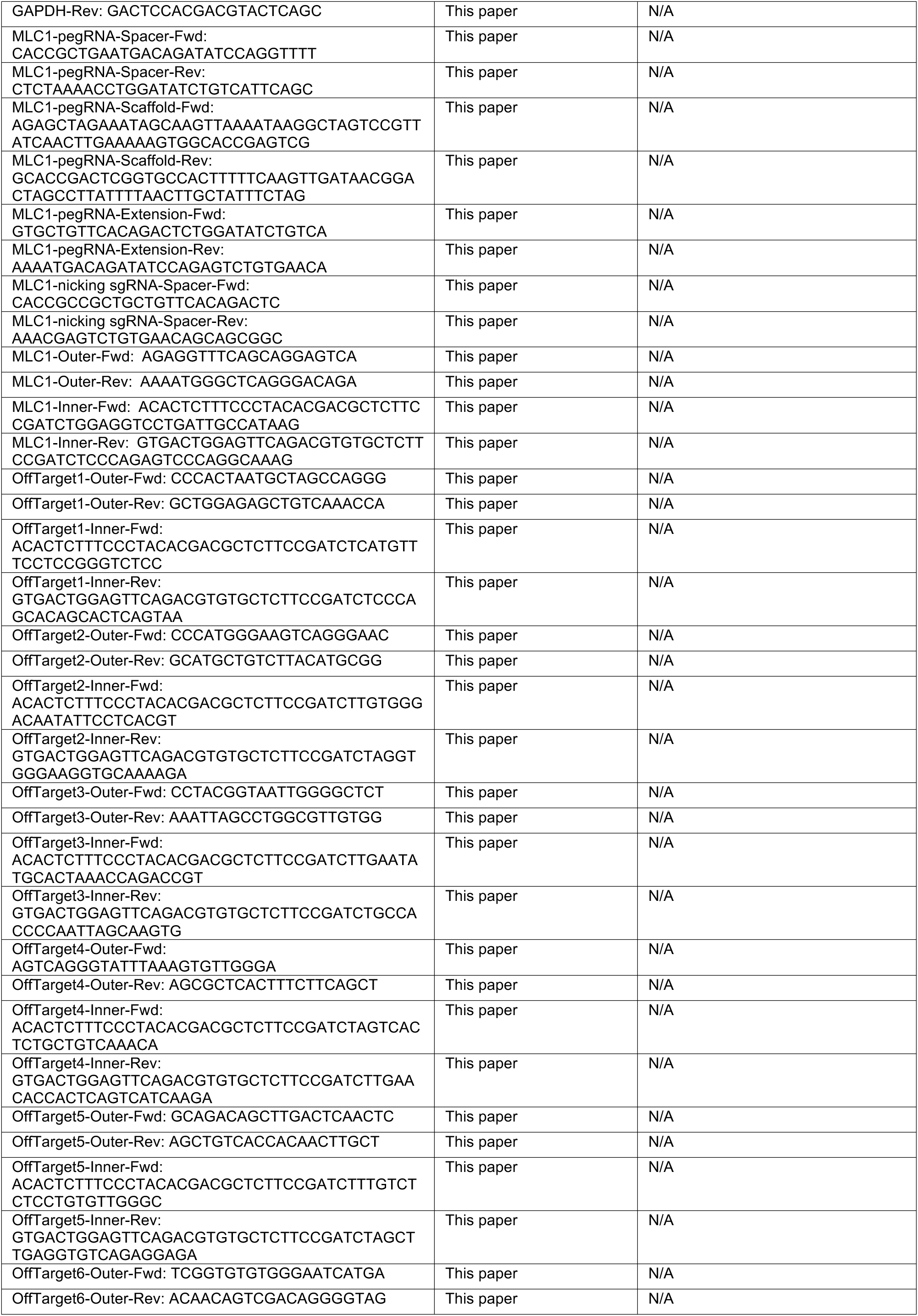
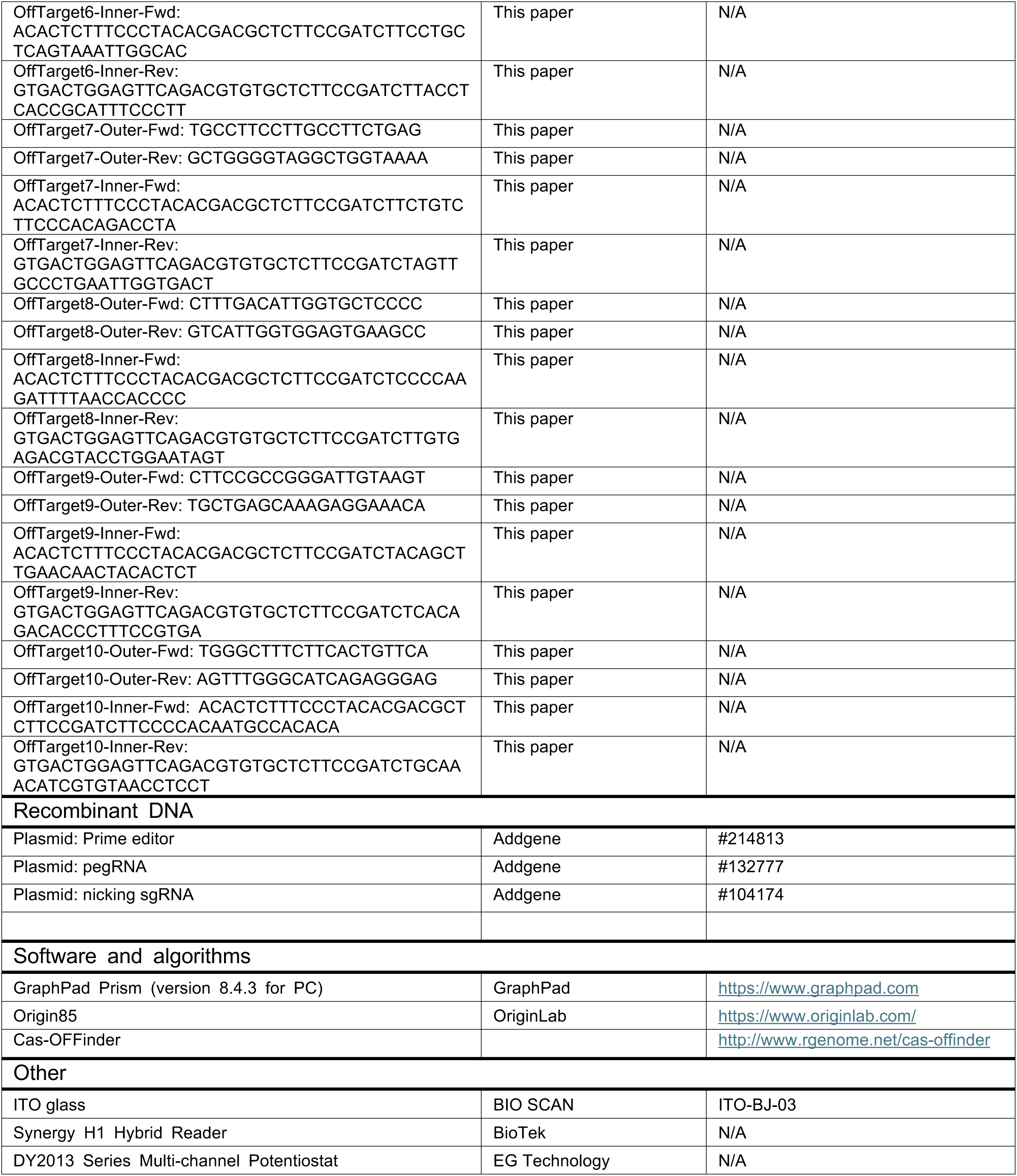

### METHOD DETAILS

#### Generation of human induced pluripotent stem cells (hiPSCs)

Human studies were performed in compliance with the Declaration of Helsinki and the study protocol was approved by the Institutional Review Board of the Catholic University of Korea (MC20TIDI0115) and Seoul National University (No. 1905-116-1035). To generate human induced pluripotent stem cells (hiPSCs), we first isolated peripheral blood mononuclear cells (PBMCs) from sex-matched controls and two MLC patients. Briefly, PBMCs were separated via centrifugation using Ficoll-Paque Plus (17-1440-03, GE Healthcare) and subsequently cultured for 5 days in StemSpan ACF (09855, Stem Cell Technology) supplemented with StemSpan CC110 (02697, Stem Cell Technology). Following the expansion period, mononuclear cells were transferred to 24-well plates pre-coated with Matrigel (1:100, 354277, Corning). For reprogramming, cells were infected with Sendai virus (CytoTune-hiPSC 2.0 Reprogramming Kit, A16518, Invitrogen) at a multiplicity of infection of three. The next day, the culture medium was replaced to Essential 8 medium (A1517001, Gibco), which was changed daily until hiPSC colonies emerged. Once colonies formed, they were manually selected and maintained in Essential medium on recombinant human vitronectin-coated plates (1:100, A25800, Gibco). On day 12 post-transduction, individual hiPSC colonies were picked and expanded for further characterization. When passaging hiPSCs using Accutase (AT-104, Innovative Cell Technologies), cells were incubated in Essential 8 medium supplanted with 10 µM ROCK inhibitor (S1049, Selleckchem) to enhance survival. Throughout the culture process, cells were maintained in a 37 °C incubator with 10% CO₂. Regular mycoplasma testing was performed throughout the process, and no contamination was detected (data now shown).

#### Characterization of control and MLC1 mutant hiPSCs

Sanger sequencing was performed to confirm the missense mutation of MLC1 gene from two MLC patients^20^. Additionally, karyotype of control and MLC1 mutant hiPSCs were analyzed using GTG-banding at 550 band resolution (GenDix). For tri-lineage differentiation, a STEMdiff Trilineage differentiation kit (05230, Stem Cell Technology) was utilized. Control and MLC1 mutant hiPSCs were cultured for 5 days (mesoderm and endoderm lineages) and 7 days (ectoderm lineage) to drive differentiation into three germ layers according to the manufacturer’s instructions. Cells were then fixed with 4% paraformaldehyde (PFA, 1.04005.1000, Merck) for 30 min and stained with the following antibodies: anti-PAX6 (1:50, sc-81649, Santa Cruz Biotechnology), anti-SM22A (1:100, sc-53932, Santa Cruz Biotechnology), and anti-FOXA2 (1:50, sc-374376, Santa Cruz Biotechnology). Finally, the expression of stem cell markers was evaluated by both quantitative PCR (qPCR) and immunocytochemistry. For qPCR, total RNA from PCMCs, control and patient hiPSCs was extracted with TRIzol reagent (9109, Takara) and cDNA was generated using a PrimeScript™ 1st strand cDNA Synthesis Kit (6110A, Takara), followed by qPCR with SYBR green (RR820, Takara) and a CFX Opus 96 Real-Time PCR system (BioRad), as previously described^32^. Quantitative analysis of mRNA expression using the double delta Ct method was carried out as previously described^33,34^. For immunocytochemistry, hiPSCs were cultured on cell culture slides (30104, SPL Life Sciences) until they were confluent. After fixation of hiPSCs with 4% PFA for 30 min, staining with the following primary antibody was performed for overnight at 4 °C: anti-OCT4 (1:50, sc-5279, Santa Cruz Biotechnology), anti-SOX2 (1:100, 630801, Biolegend), anti-NANOG (1:100, MA1-017, Invitrogen), and anti-SSEA-4 (1:100, MAB4304, Sigma-Aldrich). Then the cells were incubated with cy3-conjugated secondary antibodies (1:1000, 715-165-151 or 711-165-152, Jackson Immunoresearch) for 2 h at room temperature and visualized under a fluorescent microscope (IX73, Olympus).

#### Generation and validation of prime-edited MLC1 mutant hiPSCs

To introduce patient-relevant mutations into the MLC1 locus, control human iPSCs (∼8 × 10⁵ cells at 80% confluency) were electroporated using the 4D-Nucleofector X Unit (CA137, Lonza) with the P3 Primary Cell 3D-Nucleofector X Kit (V4XP-3012, Lonza), following the manufacturer’s protocol. Cells were transfected with a total of 1.25 μg plasmid DNA: 750 ng of a Prime Editor plasmid, 250 ng of pegRNA, and 250 ng of nicking sgRNA. Electroporated cells were seeded onto Matrigel-coated 24-well plates in StemFlex medium (A3349401, Gibco) containing 10% CloneR2 (100-0691, Stem Cell Technology) to enhance single-cell survival. Following recovery and expansion, single colonies were manually picked and clonally expanded. Each clonal cell line was lysed in 100 μL of lysis buffer (50 mM Tris-HCl (Sigma-Aldrich), 1 mM EDTA (Sigma-Aldrich), 0.005% sodium dodecyl sulfate (Sigma-Aldrich), pH 8.0 at 25 °C), supplemented with 5 μL Proteinase K (Qiagen), at 55 °C for 1 h, followed by heat inactivation at 95 °C for 10 min. Targeted deep sequencing confirmed biallelic MLC1 mutations in PCR amplicons. Clones with confirmed mutations were expanded and validated for pluripotency by immunocytochemistry and qPCR for stem cell markers. G-banding karyotype analysis confirmed genomic integrity. For off-target evaluation, potential off-target loci with up to three mismatches relative to the on-target site were identified using Cas-OFFinder (http://www.rgenome.net/cas-offinder). No loci with two or fewer mismatches were found. Among the candidates with three mismatches, ten sites were selected for targeted deep sequencing, prioritizing those with mismatches distal from the PAM for higher off-target potential. Nested PCR was used to amplify both on-target and off-target regions using KAPA HiFi HotStart polymerase (Roche), followed by a second PCR with TruSeq CD-indexed primers (Illumina) to generate NGS libraries. Libraries were purified with the QIAquick PCR purification kit (Qiagen) and sequenced on an Illumina MiniSeq or iSeq100 platform using the GenerateFASTQ workflow. Editing frequencies were determined by calculating the percentage of sequencing reads with edits relative to the total number of sequencing reads, as previously described^35^.

#### Generation of cortical organoids

We generated human cortical organoids (hCO) as previously described^21,36^. In brief, on the day of seeding (day 0), a total of 9000 iPSCs were initially treated with 20 µM ROCK inhibitor in neural induction medium, which consisted of 20% knockout serum replacement (10828028, Gibco), 1% GlutaMax (35050061, Gibco), 1% MEM-NEAA (11140035, Gibco), 0.1 mM 2-mercaptoethanol (21985023, Gibco), and 1% penicillin/streptomycin (PS-B, Capricorn) in DMEM/F12. To facilitate embryoid body (EB) formation, the cells were cultured in a U-bottom 96-well plate without surface coating. On the following day (day 1), SB-431542 (10 µM; 1615, Tocris), dorsomorphin (5 µM; P5499, Sigma), and an additional 20 µM ROCK inhibitor were added to the culture. From days 2 to 5, the medium was replaced daily with neural induction medium supplemented with 10 µM SB-431542 and 5 µM dorsomorphin. On day 6, the culture conditions were shifted to neural medium containing 20 ng/mL bFGF (100-18B, Peprotech) and 20 ng/mL EGF (AF-100-15, Peprotech), along with 2% B27 supplement without vitamin A (12587010, Gibco), 1% GlutaMax, and 1% penicillin/streptomycin in Neurobasal A medium (10888022, Gibco). This medium was refreshed daily until day 15, after which it was replaced every other day from day 17 to day 24. On day 25, each EB was transferred to an individual well of a non-coated 6-well plate, with 500 µL of neural medium supplemented with 20 ng/mL BDNF (450-02, Peprotech) and 20 ng/mL NT3 (450-03, Peprotech). The medium containing BDNF and NT3 was changed every other day until day 42. From day 43 onward, organoids were maintained in neural medium without additional factors, with medium changes occurring every 3 to 4 days. For bromo-deoxyuridine (BrdU) administration, 10 µM BrdU (B5002, Sigma) was added to the culture medium for 24 h at 30 days after hCO generation, and the cortical organoids were fixed 7 days later.

#### Histological assessments of cortical organoids

Cortical organoids were fixed at 30, 60, and 90 days after the generation in 4% PFA at 4 °C for 24 h. They were then subjected to dehydration in a 30% sucrose solution overnight at 4 °C. The organoids were embedded in an OCT compound (4583, Sakura Finetek) and rapidly frozen using liquid nitrogen. The frozen samples were then sectioned into 12 μm thick slices using a cryostat microtome (CM1850, Leica Biosystems) and subsequently mounted onto slides. After washing with 0.01 M phosphate buffered saline (PBS), tissue sections were incubated with 5% normal donkey serum and 0.3% Trion X-100 in PBS for 1 h at room temperature. Then the slides were incubated with primary antibodies overnight at 4 °C. The primary antibodies in this study were chosen based on the validation results provided by the manufacturer: anti-PAX6 (1:50, sc-81649, Santa Cruz Biotechnology), anti-SOX2 (1:100, BLG-630801, Biolegend), anti-MLC1 (1:100, ARP35249-P050, Aviva Systems Biology), mouse anti-BrdU (1:200, M0744, Dako), and anti-N-cadherin (1:1000, 33-3900, Invitrogen). Next day, after washing with 0.01 M PBS three time, slides were incubated with secondary antibodies (1:1000, 715-165-151, 711-165-152 or 706-165-148, Jackson Immunoresearch) for 2 h at room temperature. Finally, 4’6- diamidine-2’-phenylindole dihydrochloride (DAPI, 50 μg/ml, 10 236 276 001, Roche) was counterstained. For double labeling, primary antibodies to MLC1 and N-cadherin were simultaneously incubated and cy3- and Alexa Fluor 488–conjugated fluorescent secondary antibody incubations were further processed for each antibody (1:1000, 711-165-152 or 175-545-151, Jackson Immunoresearch). Finally, sections were mounted with fluorescence mounting medium (S3023, Dako) and assessed under a confocal microscope (LSM900, Carl Zeiss Microscopy).

#### Microscopic analysis and quantification

Quantitative analysis was performed using an automated image processing algorithm. We created a Python script to automatically exclude improper image areas, including tissue debris and scale bars, to select proper region of interests in hCO. This script identified the analysis targets as white mask areas **(Figure S1B)**. Other irrelevant regions, such as folded tissue areas, were manually removed from the mask **(Figure S1C)**. The analysis was conducted only on the masked areas of the tissue images. Because very weak signals can be artifactual, we set the detection threshold at 50 for each of the R, G, and B channels from the images (with R, G, and B channel values ranging from 0 to 255). The number of pixels in each color channel was measured with another Python script and overlapping areas between the color channels were also calculated. Three to six tissue sections with an interval of 60 um were assessed and the mean pixel values were presented as percentage based on the total mask area. In addition, all the quantitative analysis was performed by an observer blinded to the experimental group.

#### Western blot analysis

Cortical organoids and hiPSCs were lysed in chilled ProEX™ CETi lysis buffer (TLP-121CETi, TransLab Biosciences, Daejeon, Korea). Lysates were clarified by centrifugation, and the supernatant was collected for protein extraction. Equal amounts of protein (18–25 µg) from each sample were separated on a 10% SDS-polyacrylamide gel and transferred to a polyvinylidene difluoride membrane. The membrane was then blocked with 3% bovine serum albumin in Tris-buffered saline containing 0.05% Tween-20 for 1 h at room temperature and incubated overnight at 4°C with anti-MLC1 (1:1000, ARP35249-P050, Aviva Systems Biology), anti-glutathione peroxidase 1 (GPX1, 1:1000, LF-PA0019, Lab Frontier), anti-glutathione peroxidase 4 (GPX4, 1:1000, LF-PA0055, Lab Frontier), anti-catalase (1:1000, ab52477, Abcam), anti-superoxide dismutase 1 (SOD1, 1:500, MA1-105, Invitrogen), anti-superoxide dismutase 2 (SOD2, 1:500, sc-18503, Santa Cruz Biotechnology), OPA1 (1:1000, 612606, BD Transduction Laboratories), DRP1 (1:1000, 611112, BD Transduction Laboratories) and MFN2 (1:1000, ab56889, Abcam) antibodies. The membrane was then incubated with horseradish peroxidase-conjugated secondary antibodies (1:3,000, GTX213110-01, GTX213111-01 or AP180P, GeneTex or Millipore) at room temperature for 1 h. Bands were detected using an NICSROWEST™ ECL ADVANCE (BNROP-0009, Bionics, Seoul, Korea), after which, the blots were reprobed with mouse monoclonal anti-β-actin (1:5,000, MA1-140, Invitrogen) as a loading control. Relative quantification of the band immunoreactivity was performed by densitometric analysis using ImageQuant LAS 4000 (Fujifilm, Tokyo, Japan)^37^.

#### Biochemical assessment of mitochondrial function and oxidative stress

Mitochondrial ATP level was determined by utilizing the Mitochondrial ToxGlo assay according to the manufacturer’s procedure (G8000, Promega). Briefly, control and MLC1 mutant cortical organoids at D60 were dissociated with 0.25% trypsin for 1 h at 37 °C and resuspended in neural media. Two hundred thousand cells were then incubated with galactose-containing neural media at 37 °C for 90 min and then further incubated with ATP Detection reagent. Some of the cells were treated with oligomycin A (5 µM, 75351, Sigma) for 90 min in galactose-containing neural media for blocking mitochondrial activity. Luminescence was measured using a Victor 3 plate reader (Perkin Elmer).

The mitochondrial membrane potential was measured using MitoProbe™ TMRM Kit for Flow Cytometry (M20036, Invitrogen). One hundred thousand cells dissociated from cortical organoids were incubated with 20 nM tetramethylrhodamine methyl ester (TMRM) staining solution for 30 min at 37 °C and the fluorescent cells were inspected using a flow cytometer (FACS Aria Fusion, Beckton Dickinson) with 561 nm (excitation)/ 585 nm (emission). For some of the cells, carbonyl cyanide 3-chlorophenylhydrazone (CCCP, 50 µM) was pretreated for 5 min before TMRM treatment to uncouple the mitochondrial membrane potential serving as a positive control. The mean fluorescent intensity of TMRM was analyzed with FlowJo software (Tree Star).

Oxidative stress was assessed using 2’,7’-hichlorodihydrofluorescein diacetate (H_2_DCFDA) fluorescence. After single cell dissociation with 0.25% trypsin for 1 h, control and MLC1 mutant cells were incubated with 25 uM H_2_DCFDA (D399, Invitrogen) for 30 min in a dark incubator at 37 °C. Following incubation, fluorescence intensity was measured with a Victor 3 plate reader (Perkin Elmer) at an excitation wavelength of 488 nm and an emission wavelength of 535 nm. Some of the control and MLC1 mutant cells were treated with 2 mM hydrogen peroxide (H₂O₂) or 1 mM NAC, respectively, for 1 h at 37 °C before 30 min of H_2_DCFDA incubation.

#### Electron microscopic analysis

At 60 d after hCO generation, control and MLC1 mutant hCOs were fixed in 4% paraformaldehyde and 2.5% glutaraldehyde in 0.1M phosphate buffer (PB) for 3 hours. After rinsing in 0.1M PB, the samples were post-fixed in 1.5% potassium ferrocyanide and 1% OsO4 for 1 hr. They were then dehydrated in a graded ethanol series (50, 70, 80, 90, 95, 100%) for 10 minutes each and resin infiltration is performed with ascending concentration of resin. Mount the samples by covering resin-filled tube, and then polymerize it in a 60 °C oven for 36 hr. Polymerized resin tube containing the cells are separated from the well. Ultrathin sections were cut on an ultramicrotome (Leica Ultracut UCT) to a thickness of about 60-70 nm. The sectioned slices were collected on grids (200 mesh) and stained with 2% uranyl acetate and lead citrate. The prepared grids were examined in transmission electron microscope (HT7800, Hitachi) operating at 60 kV.

#### Fabrication of MAGO platform

To fabricate the MAGO platform, materials such as indium tin oxide (ITO) glass, gold chloride trihydrate (AuCl_3_), poly(ethylene glycol) 200 (PEG 200), Triton X-100, DPBS (Sigma-Aldrich), polydimethylsiloxane (PDMS), and plastic chambers were sourced from U.I.D, Sigma-Aldrich, and Dow Corning Corp., respectively. ITO glass substrates were prepared and cleaned by using a 1 % Triton X-100 solution, DI water, and 70 % ethanol in an ultrasonic bath. A plastic chamber was attached to the ITO electrode using PDMS, which is a bio-friendly material that allows for EC deposition of the gold mixture solution and safely cultivate the cell growth on a chip. Gold mixture solution was prepared with a 5 mM gold (III) solution with PEG 200 in a 50:1 ratio for EC detection. The gold mixture solution was deposited onto the ITO glass substrate (total area: 1.2 cm × 1.7 cm; thickness: 0.07 cm; electrical resistance: 8 ohms) was fabricated by using the multi-step potential (MSP) channel on the EC instrument (EG Technology) at 120 s, as previously reported^19^. The fabricated chips were then washed with 70% ethanol and sterilized under UV light for 1 h. Following sterilization, the gold nanostructured platform was initially coated with Matrigel (Corning) diluted 1:80 and maintained at 37°C for at least 1 h.

#### Topological and morphological characterization of the MAGO platform

To characterize the surface of the MAGO platform, we utilized FE-SEM (Carl Zeiss), AFM (Park Systems), and Conductive AFM (c-AFM, XE-100, Park Systems). FE-SEM analysis was performed at an acceleration voltage of 10 kV. Additionally, AFM analysis was conducted in tapping mode (PR-T300, Probes). For c-AFM analysis, a platinum-coated tip was used to apply an electrical potential between the tip and bottom of the MAGO. The sample bias voltage was set at 5 V. For determining the size of the gold nanoparticles and calculating the root mean square of roughness (R_q_) for each sample, the image processing and analytical program XEI was used.

#### EC monitoring of control and MLC1 mutant hCOs

To monitor control and MLC1 mutant hCOs for 75 days on the MAGO platform, the differential pulse voltammetry (DPV) experiments were conducted using a DY2013 Potentiostat (EG Technology). The fabricated Matrigel/gold nanostructure/ITO electrode acted as the working electrode, with platinum and Ag/AgCl (1 M KCl) wires serving as the counter and reference electrodes, respectively. Prior to the EC detection, the culture medium was replaced with fresh medium to eliminate potential signal interference from redox molecules and metabolites. For precise quantification, DPV signals were measured under these conditions: initial E (V) = –0.3, final E (V) = 0.5, step E (V) = 0.005, and pulse period (s) = 0.2. The DPV measurements were performed at RT. The calculated I_p_ values were analyzed by subtracting the baseline current from the current at E_p_ = –0.25 V.

#### EC assessments of metabolic activity in control and MLC1 mutant hCOs

To assess metabolic activity, Day 60–62 control and MLC1 mutant hCOs were treated with various mitochondrial inhibitors including FCCP, oligomycin (OM), rotenone, and antimycin A at concentrations of 2 µM, 4 µM, 8 µM, and 16 µM for 24 h at 37°C. For glycolytic inhibition, 2-Deoxy-D-glucose (2DG) was applied at 2.5 mM, 5 mM, or 10 mM for 24 h. After treatment, hCOs were rinsed with fresh culture media and incubated for 90 min prior to differential pulse voltammetry (DPV) measurements. Cell viability was assessed using the CellTiter-Glo® 3D Cell Viability Assay (Promega) according to the manufacturer’s instructions.

#### Statistical analysis

For statistical comparison between two groups, either a two-tailed Student’s t-test or a Mann– Whitney test was used depending on the outcome of normality assessments. Sample distributions were evaluated using the D’Agostino–Darling, Shapiro–Wilk, or Kolmogorov–Smirnov test. If at least one normality test indicated that the data were normally distributed, the Student’s t-test was applied. If all normality tests indicated non-normal distribution, the Mann–Whitney test was used. For statistical comparison of more than two groups with one independent variable, one-way ANOVA with Tukey’s post hoc test was utilized. For statistical comparison with more than two independent variables, two-way ANOVA and Tukey’s multiple comparison or two-stage linear step-up procedure of Banjamini, Krieger, and Yekutieli tests were utilized. In cases where data followed a normal distribution but the assumption of equal variances was violated, Welch’s t-test was used for comparisons between two groups, and Brown–Forsythe ANOVA followed by Dunnett’s T3 multiple comparisons test was applied for comparisons among three or more groups. Outliers were removed by ROUT’s method with Q = 1%. All statistical analyses were performed using GraphPad Prism. A *p* value of less than 0.05 was considered statistically significant. The ‘n’ values shown in figure legends represent biological replicates.

## Supplementary Information

**Figure S1.**
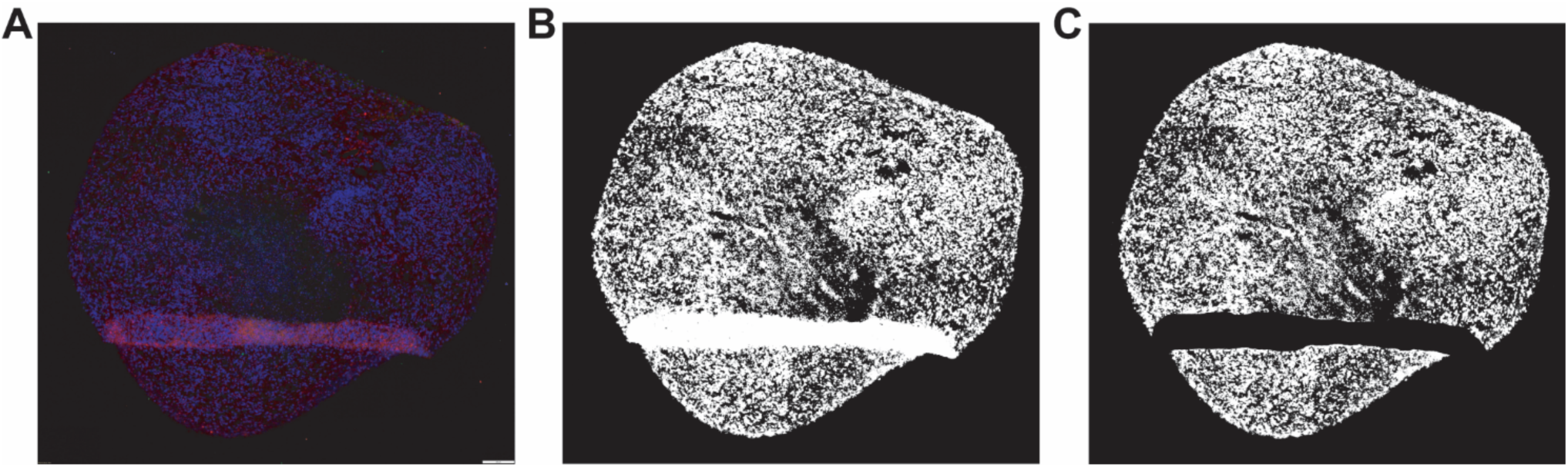
Automated image processing for quantitative analysis of human cortical organoids (hCOs). (A) Representative original image of an hCO section acquired for RGB-based image quantification. (B) Automatically generated white mask excluding debris, scale bars, and background using a custom Python script. (C) Final mask after manual removal of folded tissue areas. Quantification was performed only within masked regions using an RGB threshold of 50 (range: 0– 255).

**Figure S2.**
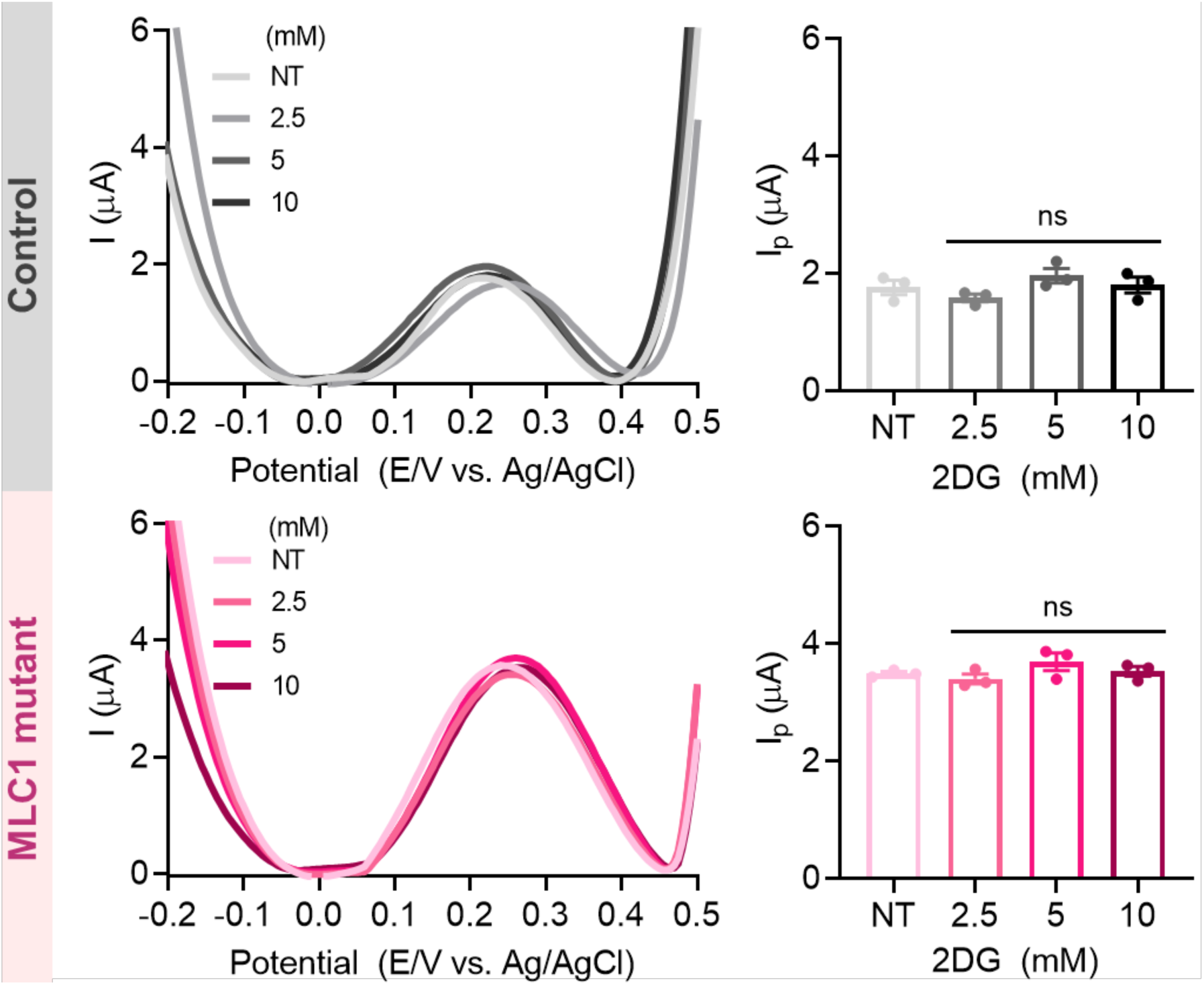
DPV graphs of hCOs (above) and MLC1 mutant hCOs (below), and the quantified DPV results are presented as bar graphs (n=3).

